# Theoretical expectations versus empirical observations in the bacterial enumeration process using serial dilution

**DOI:** 10.1101/2024.11.28.625891

**Authors:** Monika Jain, Shuhada Begum, Shuvam Bhuyan, Priya Rani Daimari, Zoomani Rabha, Chayanika Nath, Uchakankhi Kashyap, Lukapriya Dutta, Shubhra Jyoti Giri, Nishita Deka, Manabendra Mandal, Aditya Kumar, Suvendra Kumar Ray

**Affiliations:** Department of Molecular Biology & Biotechnology, Tezpur University, Tezpur–784028, Assam, India

**Author notes:** Corresponding author: Suvendra Kumar Ray.

**Keywords:** Dilution plating, CFU (Colony Forming Units), serial dilution, microliter spotting, sampling error, diluent volume

## Abstract

Considering the non-uniform distribution of bacteria in a liquid culture, accurate enumeration of bacteria by serial dilution is important yet challenging in microbiological research. The deviation of estimated population size from the actual population size is anticipated in view of the sampling error arising at dilution, which is known to increase at each successive dilution; and the sampling error associated with the sample volume, where it is inversely proportional. In the dilution process of such population following a Poisson distribution, the relation of sampling error with the sequentially diluted suspensions is yet to be empirically demonstrated. Also, as a given-fold dilution can be achieved by passaging of either a higher or a lower sample volume, the sampling error associated in this case with the volume being passaged also remains to be evaluated. In view of that, the current experimental study was undertaken to further understand bacterial distribution and the underlying sampling error in dimensions previously not demonstrated. We demonstrated that sampling error in the enumeration process is independent of the bacterial cell densities of the diluted suspensions, which was on contrary to the theoretical assumptions. Moreover, sampling error is also not dependent on the number of passages in the dilution process. Importantly, we also demonstrated that passaging either a higher or lower sample volumes during serial dilution yielded similar enumeration, which was exceptional with regards to the statistical fluctuations inherent to a Poisson distribution.

## 1. Introduction

Estimating viable microbial counts in a sample serves the fundamental pillar in microbiology and has been practiced for more than one-hundred years [1]. The most widely used method for enumeration involves sampling of the population on microbial agar plates, counting the colonies and estimating the counts in terms of Colony-Forming Units (CFU). By taking into account only the viable microbial units (capable of dividing and forming colonies), this method holds high significance in the field of clinical, food and environmental microbiology. But the high microbial abundance in any sample makes it necessary to begin with diluting it many folds before performing the sampling [2], so that each viable unit forms a distinct colony of visible size to the naked eye, which can then be counted [3]. In this method of dilution plating, serial dilution remains the classical technique being employed [2] where the desirable fold diluted suspension is attained by a series of successive dilutions with a specific dilution factor. However, for the population that follows a random or Poisson distribution like that of a bacteriological sample, the dilution although facilitates counting of the colonies upon plating; yet it is known to introduce considerable sampling error in the enumeration process which apparently increases at each successive dilution. Yet another source of sampling error is constituted by the size of the sample, where it inversely relates to the sample size [1]; such that a smaller sample volume yields greater sampling variability than a larger sample volume. This presents a natural consequence of the statistical fluctuation in the population following a random/Poisson distribution. In this context, the extent of sampling error associated with the use of different passaging volumes during serial dilution remains to be empirically demonstrated. Furthermore, the sampling error arising from progressively decreasing cell densities across the diluted suspensions also remains to be experimentally characterized. The knowledge would prove helpful in discriminating the choice of enumeration using a higher or lower dilution accordingly. Considering all these challenges of enumerating a statistically fluctuating population, the current experimental study was undertaken to further understand bacterial distribution and the underlying sampling error in dimensions previously not demonstrated. This would prove helpful in selecting the suitable approach of enumeration.

The different-fold dilutions being compared in this study for enumeration are both at a lower degree (comparison of 2-fold, 4-fold and 8-fold dilutions) and a higher degree (10-fold, 100-fold and 1000-fold dilutions). Further, the higher dilutions have been studied at two different dilution volumes: 1 mL and 10 mL. The studies were conducted using two different bacterial cell suspensions: *E. coli* DH5α and *R. pseudosolanacearum* F1C1 exhibiting two different colony morphologies, with colonies of the former being non-mucoid and latter being mucoid due to the secretion of exopolysaccharides.

## 2. Material & Methods

### 2.1. Bacterial strains and growth conditions

*E. coli* DH5α (Lab collection) was cultured in Luria Bertani (LB) (Himedia) medium (that contains 1% casein enzyme hydrolysate, 1% NaCl, 0.5% yeast extract) and incubated at 37 □C with 200 rpm shaking condition. *R. pseudosolanacearum* F1C1 (Lab collection) [4] was cultured in BG medium [5], that contains 1% peptone (Himedia), 0.1% yeast extract (Himedia) and 0.1% casamino acid (SRL), supplemented with 0.5% glucose (Himedia) and incubated at 28 □C with 150 rpm shaking condition. For preparing solid medium, 1.5% agar (Himedia) was added.

### 2.2. Culturing of strains and serial dilutions

Saturated cultures of the mentioned strains were obtained after incubating for 16 h in their respective growth conditions as mentioned above; and then serial dilutions were performed in saline (0.9% NaCl) as diluent. Three different approaches were employed for serial dilution *viz*. (i) a 10-fold, (ii) a 100-fold and (iii) a 1000-fold dilution series. The 10-, 100- and 1000-fold dilution series involves the respective series of diluting 1-part sample in 9-parts diluent, 1-part sample in 99-parts diluent and 1-part sample in 999-parts diluent. Thereby, producing the corresponding dilution ratios of 1:9, 1:99 and 1:999 respectively. These dilutions were performed for two different final volumes: 1 mL and 10 mL. The dilutions for 1 mL were performed in 1.5 mL micro-centrifuge tubes (MCT) (from Abdos) as illustrated in Figure 1 and in test tubes (from Borosil) for 10 mL dilution volume as illustrated in S.Figure 1. Tips were changed between different dilutions of the series. For a more reliable dilution, precaution was taken during taking in the sample, dispensing it on the walls of next tube in series and mixing the new suspension obtained as follows: Sample was immediately taken from a well-mixed suspension (to reduce the chances of cellular sedimentation on standby). While taking in the sample, the micro-tip was dipped just below the meniscus, not too deep to reduce bacterial adhesion on the outer surface of tip. The sample was then dispensed carefully on the walls of the next tube, without touching the diluent so that the population adhered on the outer surface of the tip would not have a chance to contact the diluent. Lastly the tip was discarded after dispensing the sample, and the sample on wall was invert-mixed after closing the MCT or covering the mouth with sterile parafilm in case of test-tube. From this suspension, next dilution of the series was similarly performed from a fresh tip.

**Figure 1.**
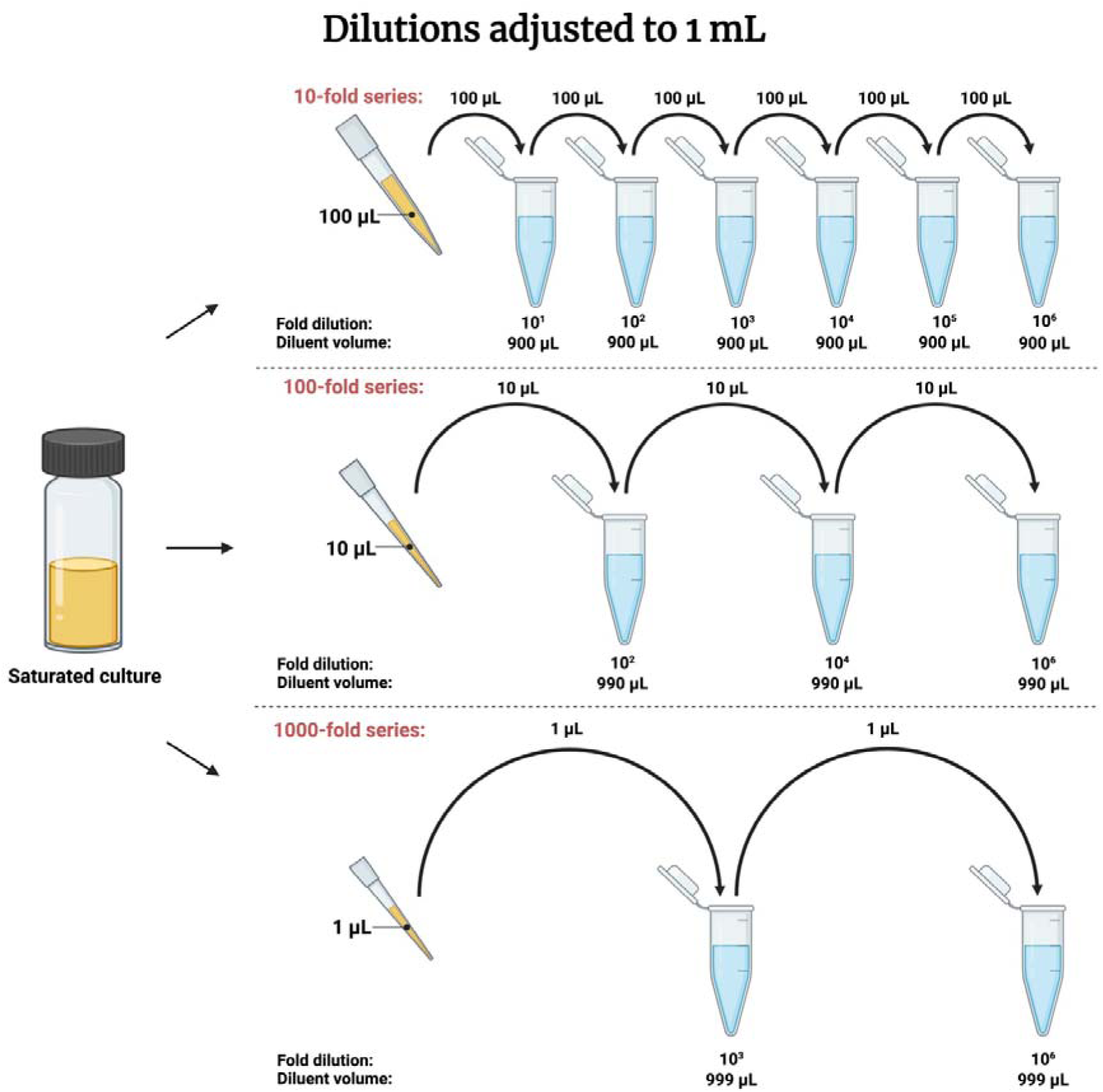
Three different ways of attaining a 10^6^ fold dilution has been shown *viz* a 10-, 100- and 1000-fold dilution series. (i) A 10-fold series (1:9 dilution ratio) involves passaging 100 μL suspension to 900 μL saline in six successive dilutions. (ii) A 100-fold series (1:99 dilution ratio) involves passaging 10 μL suspension to 990 μL saline in three successive dilutions. (iii) A 1000-fold series (1:999 dilution ratio) involves passaging 1 μL suspension to 999 μL saline in two successive dilutions. Bacterial culture is shown in yellow in the screw caped bottle. Diluent (saline) is shown in blue in micro-centrifuge tubes. Created with Biorender.com.

#### 2.2.1. Dilutions to a final volume of 1 mL

As illustrated in Figure 1, for a 1 mL final volume, a 10-fold dilution series was performed by adding 100 µL of culture to 900 µL of saline in the first step to obtain a 10^1^-fold dilution, subsequently 100 µL of this suspension was added to the next tube in the series containing 900 µL of saline to obtain a 10^2^-fold dilution cell suspension. This was repeated until a cell suspension of 10^7^-fold dilution was obtained. Similarly, a 100-fold dilution series was performed by adding 10 µL of culture to 990 µL of saline in the first step to obtain a 10^2^-fold dilution. In subsequent steps, 10^4^- and 10^6^-fold dilutions were similarly obtained. Lastly, to proceed with a 1000-fold dilution series, 1 µL of culture was added to 999 µL of saline to obtain 10^3^-fold dilution in first step and 10^6^-fold dilution in second step. In both the latter dilution series (100-fold and 1000-fold series), the final 10^7^-fold dilution was obtained by performing a 10-fold dilution from their respective 10^6^-fold diluted suspensions.

#### 2.2.2. Dilutions to a final volume of 10 mL

Dilutions were performed in a similar way as mentioned above, only the volumes of sample and diluent taken were both up scaled (although same in proportion) for adjusting it to 10 mL volume as illustrated in S.Figure 1. In a 10-fold dilution series 1 mL of culture was added to 9 mL of saline to obtain a 10^1^-fold dilution, which was repeated with subsequent diluents in series to obtain 10^7^-fold dilution. In a 100-fold dilution series, 100 µL of culture was added to 9.900 mL of saline to obtain a 10^2^-fold dilution. In subsequent steps, 10^4^- and 10^6^-fold dilutions were similarly obtained. Lastly, in a 1000-fold dilution series, 10 µL of culture was added to 9.990 mL of saline to obtain 10^3^-fold dilution and then 10^6^-fold dilution in second step. In both the latter dilution series, a 10^7^-fold dilution was obtained by performing a 10-fold dilution from their respective 10^6^-fold diluted suspensions.

### 2.3. Plating and colony observation

Plating was done from a 10^6^- and 10^7^-fold diluted suspensions obtained from each of the series for *E. coli* DH5α and at 10^7^- and 10^8^-fold diluted suspensions obtained from each of the series for *R. pseudosolanacearum* F1C1. A higher fold dilution was chosen for *R. pseudosolanacearum* F1C1 as (i) with the saturated cultures of the two, the number of colonies observed in this case were always higher than that of *E. coli* DH5α by 2-6 times at same fold dilution (this can also be observed in Table 1) and (ii) *R. pseudosolanacearum* forms typical mucoidal colonies once they appear completely in plate; also they show twitching motility over solid media unlike *E. coli* [6,7]; the latter one forming non-mucoidal colonies. So apparently, colonies of the former one had greater tendency to merge with other such colonies.

**Table 1.** Colony numbers obtained in different plating volumes of the two different bacterial cell suspensions.

| Colony count data of 10 spots |  |  |  |  |  |  |  |  |
| --- | --- | --- | --- | --- | --- | --- | --- | --- |
| Bacterial samples | *E. coli* DH5α | | | | *R. pseudosolanacearum* F1C1 | | | |
| Dilution factors | $10^6$ | | | | $10^7$ | | | |
| Spotting volume | 5 μL | 10 μL | 15 μL | 20 μL | 5 μL | 10 μL | 15 μL | 20 μL |
|  | 10 | 27 | 47 | 44 | 7 | 6 | 11 | 14 |
|  | 11 | 24 | 42 | 40 | 4 | 6 | 10 | 16 |
|  | 12 | 21 | 36 | 49 | 1 | 10 | 17 | 15 |
|  | 15 | 28 | 39 | 45 | 4 | 8 | 11 | 12 |
|  | 12 | 26 | 36 | 42 | 5 | 6 | 9 | 17 |
|  | 13 | 24 | 34 | 44 | 2 | 6 | 11 | 15 |
|  | 12 | 28 | 42 | 45 | 4 | 8 | 9 | 19 |
|  | 18 | 19 | 39 | 49 | 5 | 14 | 12 | 11 |
|  | 15 | 25 | 38 | 41 | 4 | 4 | 6 | 15 |
|  | 14 | 24 | 36 | 42 | 6 | 8 | 10 | 19 |
| Mean | 13.2 | 24.6 | 38.9 | 44.1 | 4.20 | 7.60 | 10.60 | 15.30 |
| SD | 2.35 | 2.91 | 3.87 | 3.07 | 1.75 | 2.80 | 2.80 | 2.63 |
| CoV | 0.18 | 0.12 | 0.10 | 0.07 | 0.42 | 0.37 | 0.26 | 0.17 |
| CFU estimate | $2.6 \times 10^9$ | $2.5 \times 10^9$ | $2.6 \times 10^9$ | $2.2 \times 10^9$ | $8.4 \times 10^9$ | $7.6 \times 10^9$ | $7.1 \times 10^9$ | $7.7 \times 10^9$ |

Standard deviation (SD), Coefficient of Variation (CoV) and CFU estimate calculated for parent culture in each of these cases are represented.

For plating, microliter spotting approach was followed [8]. Colonies were observed and counted after 12-14 h of incubation in case of *E. coli* DH5α and after 48 h of incubation in case of *R. pseudosolanacearum* F1C1. Although with the latter one, colonies appear fully after 48 h of incubation (due to a slower generation time), but the simultaneous production of copious amounts of EPS by the bacteria makes it fluidal and difficult to count at this point. So, in this case, a prior observation of the colonies was also made at 28 h post incubation, where tiny colonies although not so prominent can be lightly visualized against light and accordingly counted. These were tallied after the colonies completely appear.

### 2.4. Study on the influence of plating volume on bacterial enumeration

To study the influence of plating/sampling volumes on bacterial enumeration, ten different spots of each of the different volumes: 5 µL, 10 µL, 15 µL and 20 µL were made in separate nutrient agar plates. These were sampled from a specific suspension which was at 10^6^ fold dilution for *E. coli* DH5α and at 10^7^ fold dilution for *R. pseudosolanacearum* F1C1, obtained simply from a 10-fold series performed for 1 mL final volume. Plates were then kept under incubation for colony observation. After incubation, average number of colonies per spot was calculated in each case.

### 2.5. Study on the influence of passaging volume on bacterial enumeration

To study the influence of passaging volume used in serial dilution on bacterial enumeration, all the three dilution series (10-fold, 100-fold and 1000-fold series) were performed for the two final volumes: 1 mL and 10 mL in the independent experiments generated from different cultures. For plating, ten spots of 10 µL each were made from a 10^6^- and 10^7^-fold diluted suspensions of *E. coli* DH5α and a 10^7^- and 10^8^-fold diluted suspensions of *R. pseudosolanacearum* F1C1. Plates were then kept under incubation for colony observation. After incubation, average number of colonies per spot was calculated in each case.

### 2.6. Study on the influence of dilution volume on bacterial enumeration

For this study, all the dilution series were performed for both 1 mL and 10 mL dilution volume and were generated from same culture, so that except the factor of dilution volume chosen, all other factors remain constant. This study was performed with *E. coli* DH5α as representative. Plating was done from a 10^6^ and 10^7^ fold diluted suspension. For each of the three dilution series, ten spots of 10 µL each were made for it being performed at 1 mL and 10 mL final volume in the different halves of same plate to enable clear comparison. The plates were then kept under incubation and later average number of colonies per spot was calculated for all the cases.

### 2.7. Study on the relation of sampling error with bacterial cell-density

First, a 10^6^-fold diluted suspension was obtained normally *via* a 10-fold dilution series performed for 2 mL dilution volume. It involved passaging 200 µL sample volume in 1800 µL saline. Later from this 10^6^-fold diluted suspension, a single-step 2-fold, 4-fold and 8-fold dilutions were independently performed (as depicted in S.Figure 2(a)) for 1 mL dilution volume, by the respective passaging of 500 µL, 250 µL and 125 µL sample volumes in the corresponding 500 µL, 750 µL and 875 µL saline. For plating, twenty spots of 20 µLeach were spotted from each of the 2-fold, 4-fold and 8-fold diluted suspension. Likewise, three trials were performed from the same primary culture. This study was performed with *E. coli* DH5α as representative. The plates were then kept under incubation and later, average number of colonies, Standard Deviation (SD) and Coefficient of Variation (CoV) were calculated for each of the three cases.

### 2.8. Study on the influence of number of dilution passages on bacterial enumeration

Apart from the single-step obtained 4-fold and 8-fold diluted suspension as mentioned above, the 4-fold and 8-fold diluted suspensions were also generated *via* the 2-fold series in the successive two and three steps respectively (as depicted in S.Figure (2b)). Twenty spots of 20 µL each were spotted from each of these suspensions in addition to the twenty spots of the single-step generated 4-fold and 8-fold suspensions. Three trials were likewise performed from the same primary culture. The plates were then kept under incubation and later, average number of colonies, SD and CoV were calculated for all the cases.

### 2.9. Statistical analyses

#### 2.9.1. Coefficient of Variation (CoV)

For comparing different sample volumes used to estimate bacterial cell density, CoV (which is a measure of variability in a population) was calculated among the colony numbers observed in the ten replicates of each sampling volumes used. First, average number of colonies and standard deviation (SD) were calculated for each of the different sample volumes, then CoV was calculated by dividing standard deviation by average.

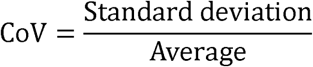

#### 2.9.2. Correlation analysis

To study the correlation between different sampling volumes used for bacterial cell density estimation and CoV obtained in the colony count data corresponding to each sampling volume, Pearson’s correlation analysis was performed. The Pearson’s correlation coefficient (r) was calculated as:

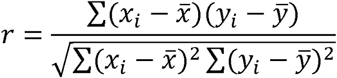

where x_i_ = values of x-variable (here, x takes the values of different sample volumes); x = mean of the values of x-variable; y_i_ = values of y-variable (here, y takes the values of CoV corresponding to each values of x); y = mean of the values of y-variable

#### 2.9.3. One-way ANOVA

For comparing the mean of sample population of the suspensions obtained from different dilution series, one-way ANOVA test was performed in MS Excel at α=0.05 with H_0_ stating that there is no significance difference between means of sample population of suspensions obtained from these different dilution series and H_1_ stating that the means of sample population of suspensions obtained from these dilution series are significantly different.

#### 2.9.4. Bonferroni post-hoc test

In cases where one-way ANOVA proved significant difference between the groups, a post-hoc test was performed using Bonferroni corrected p-value to identify which group differ significantly. This was calculated as:

Bonferroni-corrected p-value = α / number of comparative tests performed

When p-value of comparative tests < Bonferroni-corrected p-value, the test is significant.

#### 2.9.5. Mann-Whitney U test

A two-tailed Mann-Whitney U test was performed to compare the mean difference of colony counts obtained with dilutions performed for 1 mL and 10 mL volumes. It was also performed to compare the mean difference of colony counts obtained with the 4-fold and 8-fold diluted suspensions when attained in a single step versus when attained in two and three respective steps *via* the 2-fold dilution series. Test is significant when U_Stat <_ U_Crit_. U_Stat_ = minimum of the two values, U_1_ and U_2_.

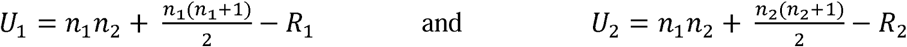

Where, *n_1_*and *n_2_* represents sample size of the two groups under comparison; *R_1_* and *R_2_* denotes rank sum of the two groups respectively when all of them are arranged in ascending order.

## 3. Result

### 3.1. A higher sample volume of a bacterial suspension is a better representation of the population size than a lower sample volume

During bacterial enumeration by dilution method, a specific volume of sample is stepwise passaged to a specific volume of the diluent. As distribution of bacteria is random in a culture, there would be anomalies associated with the colony numbers being passaged. In consideration of the random distribution, the sample volume being passaged is likely to play an important role in the enumeration. To get an insight into the influence of sample volume on consistency of bacterial count, we studied the variation prevalent in colony count in the plating of different sample volumes: 5 μL, 10 μL, 15 μL and 20 μL of a diluted bacterial suspension. The colony numbers obtained in 10 spots of each of these sample volumes is represented in Table 1. To enable clear distinction of colonies in spotted area, the suspension used was at 10^6^-fold dilution of the saturated culture in case of *E. coli* and at 10^7^-fold dilution in case of *R. pseudosolanacearum*. In case of *E. coli*, the CoV observed in colony numbers obtained with 5 μL, 10 μL, 15 μL and 20 μL spotting were 0.18, 0.12, 0.10 and 0.07 respectively. When the same study was conducted with *R. pseudosolanacearum*, the CoV was found to be 0.42, 0.37, 0.26 and 0.17 respectively for 5 μL, 10 μL, 15 μL and 20 μL spotting. Clearly, for both the independently tested bacterial strains, the coefficient of variation (CoV) of bacterial count in the different spots made of a given sampling volume was observed to be highest with the lowest sampling volume used (5 μL) and lowest with the highest sampling volume used (20 μL). Similar observations were recorded in four repetitive experiments (S.Table1). This is in concordance with the view of random distribution of bacteria in a culture; as with a randomly distributed population, a larger sampling size is supposed to give more consistency in density estimation. As expected, a strong negative correlation was found between sampling volumes of the suspension used and coefficient of variation (CoV) of bacteria count, with the same trend being observed for both the bacterial species (r = −0.97 for *E. coli* and r = −0.99 for *R. pseudosolanacearum*) (Figure 2). This suggests that higher the sampling volume of the suspension, greater the consistency in bacterial cell count and lesser the variability. Hence, higher sampling volume is a better representation of the bacterial population.

**Figure 2.**
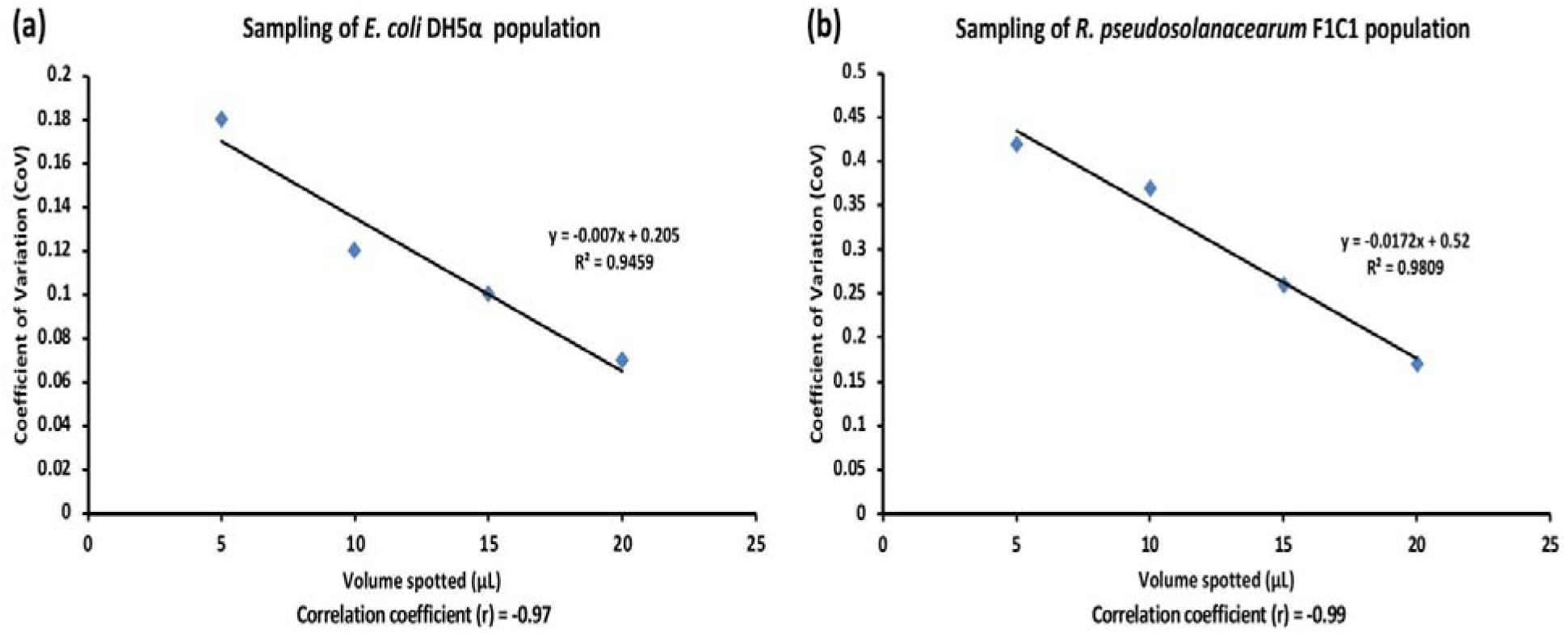
Correlation between suspension volume spotted (X-axis) and coefficient of variation (CoV) obtained among the colony numbers (Y-axis) of (a) *E. coli* and (b) *R. pseudosolanacearum*. Pearson’s correlation coefficient (r) is represented for each case. A strong negative correlation has been observed in each case

### 3.2. Stepwise passaging of 1 mL, 100 μL and 10 μL (but not 1 μL) are similar with regards to CFU estimation of E. coli DH5α

In theory, to make a 10^6^-fold dilution of an original culture different dilution series such as ten or hundred or thousand-fold can be used which has been shown in Figure 1. As evident from the Figure, in case of a higher fold dilution series such as thousand, only 1 μL of the bacterial suspension is passaged twice to achieve 10^6^-fold dilution whereas in case of a lower fold dilution series such as ten, 100 μL of bacterial suspension is passaged six times. In the intermediary hundred-fold dilution series, 10 μL of bacterial suspension is passaged thrice. Although, the 10-fold dilution series is widely practiced by the researchers considering the advantage of larger volume passaging in view of the random distribution of bacteria, but its effectiveness over the 100- and 1000-fold series is yet to be demonstrated. To compare the effectiveness of these dilution series, we estimated CFU of *E. coli* culture using these three-dilution series *viz* a 10-fold, a 100-fold and a 1000-fold dilution series as discussed below.

The CFU of bacterial suspension obtained from the three different dilution series (a 10-, 100-, and 1000-fold dilution series) is represented in Table 2(a) and a representative result is shown in Figure 3(a). The 10 μL spotting of 10^6^-fold dilution obtained from a 10-fold and a 100-fold dilution series produced average 11.9 ± 4.1 and 11.3 ± 4.5 numbers of colonies per spot respectively; however, that obtained from a 1000-fold dilution series produced average 28.6 ± 1.8 numbers of colonies per spot. Similarly with 10^7^ fold dilution average number of colonies per spot obtained from a 10- and 100-fold dilution series were 1.4 ± 1.0 and 1.1 ± 1.2 respectively; however a 1000-fold dilution series produced average 3.5 ± 1.4 number of colonies per spot. A p-value < 0.05 was indicative of significant differences among the groups, and thus a post-hoc test was followed in each case to identify the group being significantly different. This was observed to be the 1000-fold series in almost all of these cases. The results indicated that a 10-fold and 100-fold dilution series are producing similar CFU count whereas the 1000-fold dilution series is giving a higher CFU count than the other two. This trend was further observed in all the three repetitive experiments (Table 2(a)). In addition the CFU count by the 1000-fold dilution series was inconsistent. This was perplexing for us to find out if a 1000-fold dilution series is more accurate than the other two dilution series. As the results by 1000-fold dilution series were inconsistent, we thought pipetting a smaller volume like 1 μL might be a more complex procedure while taking in as well as dispensing out the suspension. To avoid any bias related to pipette, we repeated the experiments using pipettes from three neighboring labs for pipetting 1 μL volume. In this case also, an inconsistently higher CFU count was observed with the 1000-fold series. This led us to assume that 1 μL pipetting indeed is more complex and prone to inconsistencies than 10 μL and 100 μL pipetting. This might be because a minor deviation in either taking in and/or dispensing out the 1 μL suspension would contribute a larger percentage of change in the total volume.

**Figure 3.**
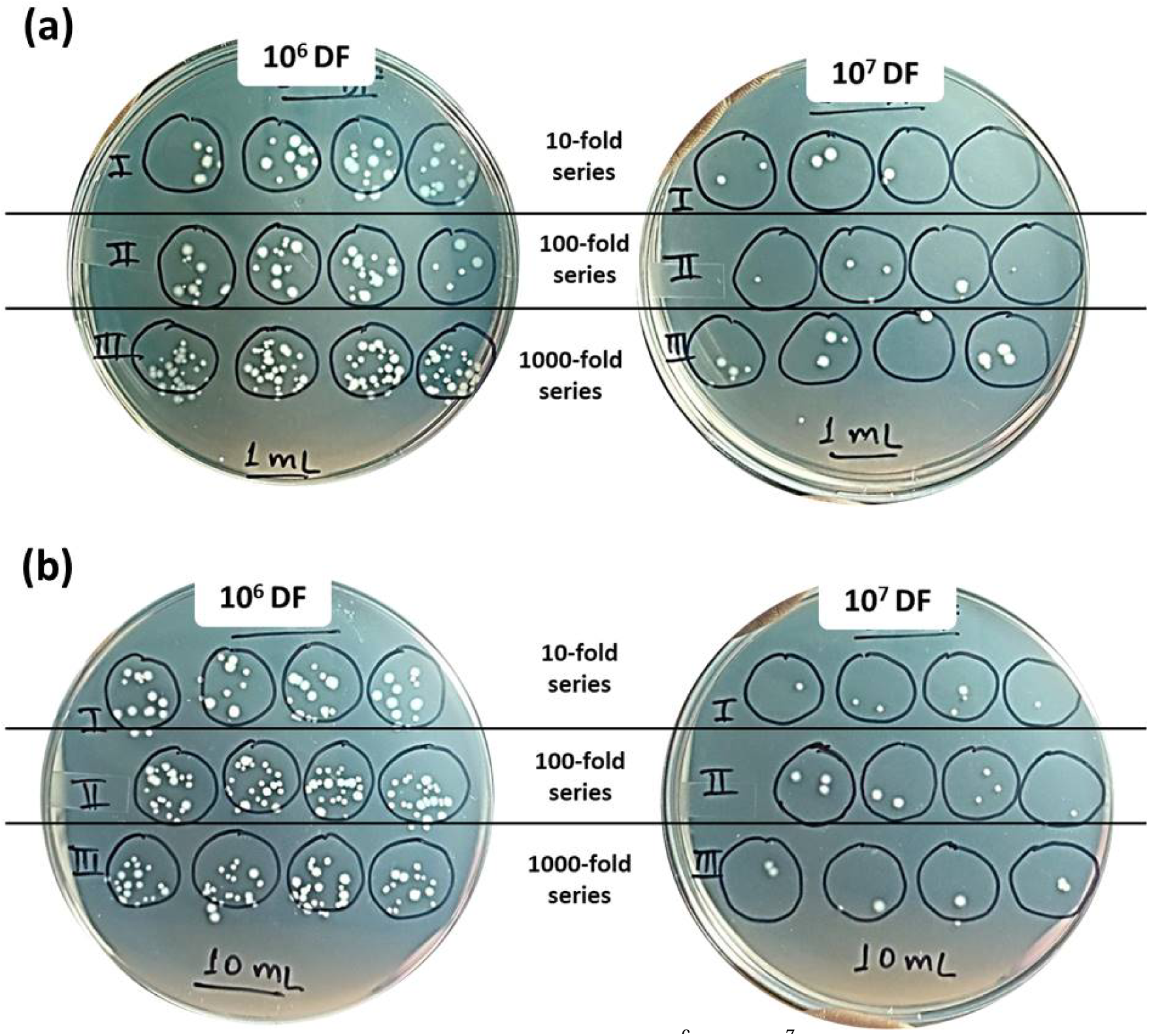
*E. coli* colonies at selected fold dilutions (10^6^ and 10^7^) obtained with the different dilution series. These series were performed first to a final volume of (a) 1 mL and later to a final volume of (b) 10 mL from independent cultures. At 1 mL dilution volume, a clearly greater number of colonies were observed with the 1000-fold series; however, at 10 mL dilution volume, similar numbers of colonies were observed with all the three dilution series

**Table 2(a).** Average colony numbers (along with standard deviations) of *E. coli* DH5α obtained from different dilution series performed to a final 1 mL volume.

| Dilutions adjusted to a total 1 mL volume (*E. coli* DH5α) |  |  |  |  |  |  |  |
| --- | --- | --- | --- | --- | --- | --- | --- |
| Dilution factors |  | 10<sup>6</sup> |  |  | 10<sup>7</sup> |  |  |
| Dilution series |  | 10 fold | 100 fold | 1000 fold | 10 fold | 100 fold | 1000 fold |
| Dilution ratios |  | 1:9<br>(A) | 1:99<br>(B) | 1:999<br>(C) | 1:9<br>(A) | 1:99<br>(B) | 1:999<br>(C) |
| Experiment 1 | Colony Count | 11.9 ± 4.1 | 11.3 ± 4.5 | 28.6 ± 1.8 | 1.4 ± 1.0 | 1.1 ± 1.2 | 3.5 ± 1.4 |
|  | One-way ANOVA | F (2, 27) = 71.64, p = 1.6 x 10<sup>-11</sup> |  |  | F (2, 27) = 11.60, p = 2.3 x 10<sup>-4</sup> |  |  |
|  | Bonferroni post-hoc test | (C) > (A), (B); (A) □ (B) |  |  | (C) > (A), (B); (A) □ (B) |  |  |
| Experiment 2 | Colony Count | 9.5 ± 1.6 | 8.3 ± 3.4 | 28.8 ± 5.2 | 1.8 ± 1.2 | 1.8 ± 1.3 | 3.5 ± 2.0 |
|  | One-way ANOVA | F (2, 27) = 108.21, p = 1.3 x 10<sup>-13</sup> |  |  | F (2, 27) = 4.08, p = 2.8 x 10<sup>-2</sup> |  |  |
|  | Bonferroni post-hoc test | (C) > (A), (B); (A) □ (B) |  |  | (A) □ (B) □ (C) |  |  |
| Experiment 3 | Colony Count | 22.5 ± 3.3 | 19.9 ± 3.5 | 45.4 ± 5.1 | 3.4 ± 1.8 | 2.9 ± 1.6 | 7.5 ± 2.0 |
|  | One-way ANOVA | F (2, 27) = 120.33, p = 3.5 x 10<sup>-14</sup> |  |  | F (2, 27) = 19.15, p = 6.6 x 10<sup>-6</sup> |  |  |
|  | Bonferroni post-hoc test | (C) > (A), (B); (A) □ (B) |  |  | (C) > (A), (B); (A) □ (B) |  |  |

F-statistic and p-value calculated from One-way ANOVA and interpretation of Bonferroni post-hoc tests for the different comparable groups are represented.

To compare the accuracies of these dilution series after eliminating the complex factor of 1 μL pipetting, all the three series were re-performed this time to a final 10 mL volume by passaging stepwise 1 mL, 100 μL and 10 μL respectively in the 10-, 100- and 1000-fold dilution series. The CFU now obtained with each of these dilution series are represented in Table 2(b) and a representative result is shown in Figure 3(b). The 10 μL spotting of 10^6^ fold dilution obtained from a 10-, 100- and 1000-fold dilution series produced average 13.7 ± 4.2, 14.7 ± 2.1 and 16.7 ± 3.9 numbers of colonies per spot respectively. Similarly with the 10^7^ - fold dilution, average number of colonies obtained from the 10-, 100- and 1000-fold dilution series were 1.6 ± 1.4, 2.6 ± 1.8 and 2.0 ± 0.9 respectively. This trend was further observed in all the three repetitive experiments. It was interesting to observe this time a similar number of colonies with all the three dilution series. A p-value > 0.05 as obtained in one-way ANOVA was indicative of no significant differences between the colony numbers obtained from the different dilution series. This proves a 1000-fold dilution series to be equally accurate as the 10-fold and 100-fold dilution series provided that pipetting of 1 μL is not carried out.

**Table 2(b).**
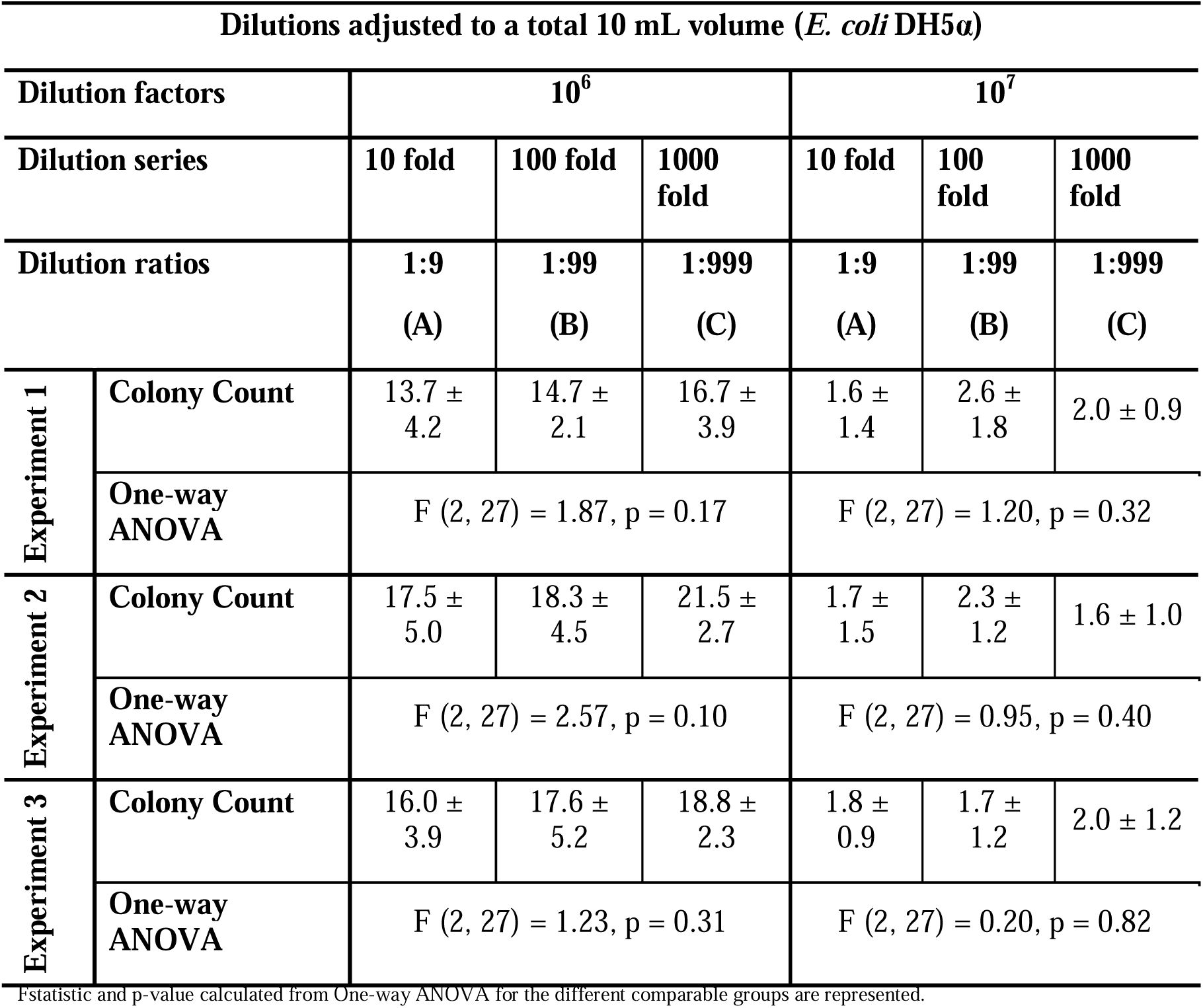
Average colony numbers (along with standard deviations) of *E. coli* DH5α obtained from different dilution series performed to a final 10 mL volume.

| Dilutions adjusted to a total 10 mL volume (*E. coli* DH5α) | | | | | | | |
| --- | --- | --- | --- | --- | --- | --- | --- |
| Dilution factors |  | 10<sup>6</sup> |  |  | 10<sup>7</sup> |  |  |
| Dilution series |  | 10 fold | 100 fold | 1000 fold | 10 fold | 100 fold | 1000 fold |
| Dilution ratios |  | 1:9<br>(A) | 1:99<br>(B) | 1:999<br>(C) | 1:9<br>(A) | 1:99<br>(B) | 1:999<br>(C) |
| Experiment 1 | Colony Count | 13.7 ± 4.2 | 14.7 ± 2.1 | 16.7 ± 3.9 | 1.6 ± 1.4 | 2.6 ± 1.8 | 2.0 ± 0.9 |
|  | One-way ANOVA | F (2, 27) = 1.87, p = 0.17 |  |  | F (2, 27) = 1.20, p = 0.32 |  |  |
| Experiment 2 | Colony Count | 17.5 ± 5.0 | 18.3 ± 4.5 | 21.5 ± 2.7 | 1.7 ± 1.5 | 2.3 ± 1.2 | 1.6 ± 1.0 |
|  | One-way ANOVA | F (2, 27) = 2.57, p = 0.10 |  |  | F (2, 27) = 0.95, p = 0.40 |  |  |
| Experiment 3 | Colony Count | 16.0 ± 3.9 | 17.6 ± 5.2 | 18.8 ± 2.3 | 1.8 ± 0.9 | 1.7 ± 1.2 | 2.0 ± 1.2 |
|  | One-way ANOVA | F (2, 27) = 1.23, p = 0.31 |  |  | F (2, 27) = 0.20, p = 0.82 |  |  |

F-statistic and p-value calculated from One-way ANOVA for the different comparable groups are represented.

### 3.3. Stepwise passaging of 1 mL, 100 μL and 10 μL (but not 1 μL) are similar with regards to CFU estimation of R. pseudosolanacearum F1C1

We further performed this CFU estimation study for a different bacterium: *R. pseudosolanacearum*, a phytopathogen, which in contrast to *E. coli*, forms mucoidal colonies on plate due to the production of exopolysaccharides [7,9]. This study was undertaken to know whether the accuracies observed previously with the different dilution series holds true for the other bacterial suspension too. Unlike *E. coli*, a higher fold dilution (10^7^ and 10^8^) was chosen for plating this time as the mucoidal nature of the colonies and twitching motility exhibited by *R. pseudosolanacearum* over solid media [6] leads towards a greater tendency for the colonies to merge among each other. Also the CFU/mL of the saturated culture of *R. pseudosolanacearum* was always observed to be higher than that of *E. coli* by 2-6 times (this can also be observed in Table1). So plating a higher fold dilution in this case will enable clear distinction of the colonies. When the dilutions were performed to a final volume of 1 mL, average number of colonies observed per 10 μL spot of 10^7^ fold dilution were 6.0 ± 2.5, 5.4 ± 1.0 and 14.1 ± 3.5 respectively with a 10-, 100- and 1000-fold dilution series (Table 3(a)). Similarly with 10^8^ fold dilution, average number of colonies per spot obtained from a 10-, 100-, and 1000-fold dilution series were 0.6 ± 0.8, 0.4 ± 0.7 and 2.5 ± 1.3 respectively. A p-value < 0.05 was indicative of significant differences among the groups, and thus a post-hoc test was followed in each case to identify the group being significantly different. This was observed to be the 1000-fold series in almost all of these cases. A representative result is shown in Figure 4(a). Aligning with our previous observation, CFU recorded with the 1000-fold series were again observed to be significantly higher and were inconsistent than that recorded with the other two dilution series. This further highlight the potential complexities associated with 1 μL pipetting in the 1000-fold dilution series and so to eliminate it, the dilutions were re-performed to a final volume of 10 mL as done previously with *E. coli* suspension. In this case, average number of colonies observed per 10 μL spot of 10^7^ fold dilution were 9.8 ± 1.9, 8.4 ± 4.3 and 7.2 ± 2.0 respectively with a 10-, 100- and 1000-fold dilution series (Table 3(b)). Similarly with 10^8^ fold dilution, average number of colonies per spot obtained from a 10-, 100-, and 1000-fold dilution series were 0.3 ± 0.7, 0.5 ± 0.7 and 0.2 ± 0.4 respectively. It was interesting to note that the colony numbers observed were similar in this case and this trend was repetitive in all the three experiments. A representative result is shown in Figure 4(b). A p-value > 0.05 as obtained in one-way ANOVA was indicative of no significant differences between the colony numbers obtained from the different dilution series. This was again in concordance with our previous observation of *E. coli* CFU estimation. This establishes 1000-fold dilution series to be equally accurate as the 10-fold and 100-fold dilution series when 1 μL pipetting is not carried out and validates our understanding on the inconsistencies and complexities associated with 1 μL pipetting. In summary, when performing the dilutions to a final volume of 1 mL, a 10- and 100-fold series proved to be effective for CFU estimation and when performing them to a final volume of 10 mL, all the 10-, 100- and 1000-fold dilution series proved to be equally effective for CFU estimation. But considering the time, cost and labor efficiency of the approach with no compromise on accuracy of CFU estimation, a 1000-fold dilution series shall be preferred when performing the dilutions for 10 mL final volume and a 100-fold dilution series shall be preferred when performing the dilutions for 1 mL final volume. This is because a relatively lower number of passages are involved in these processes compared to the 10-fold series.

**Figure 4.**
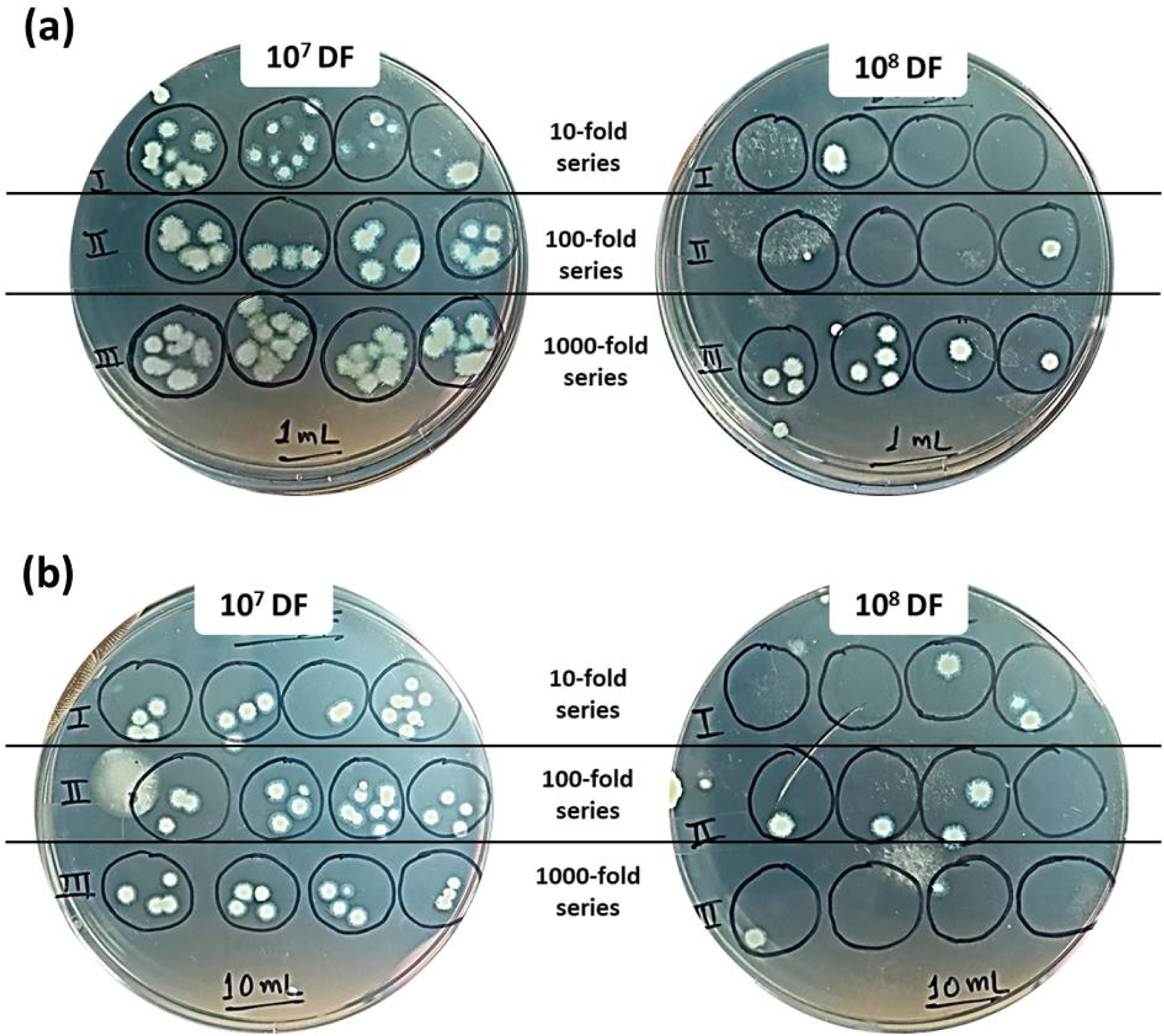
*R. pseudosolanacearum* colonies at selected fold dilutions (10^7^ and 10^8^) obtained with the different dilution series. These series were performed first to a final volume of (a) 1 mL and later to a final volume of (b) 10 mL from independent cultures. At 1 mL final volume, a clearly greater number of colonies were observed with the 1000-fold series; however, at 10 mL final volume, similar numbers of colonies were observed with all the three dilution series

**Table 3(a).** Average colony numbers (along with standard deviations) of *R. pseudosolanacearum* F1C1 obtained from different dilution series performed to a final 1 mL volume.

| Dilutions adjusted to a total 1 mL volume (*R. pseudosolanacearum* F1C1) |  |  |  |  |  |  |  |
| --- | --- | --- | --- | --- | --- | --- | --- |
| Dilution factors | | $10^7$ | | | $10^8$ | | |
| Dilution series |  | 10-fold | 100-fold | 1000-fold | 10-fold | 100-fold | 1000-fold |
| Dilution ratios |  | 1:9<br>(A) | 1:99<br>(B) | 1:999<br>(C) | 1:9<br>(A) | 1:99<br>(B) | 1:999<br>(C) |
| Experiment 1 | Colony Count | 6.0 ± 2.5 | 5.4 ± 1.0 | 14.1 ± 3.5 | 0.6 ± 0.8 | 0.4 ± 0.7 | 2.5 ± 1.3 |
|  | One-way ANOVA | F (2, 27) = 35.95, p = 2.4 x 10<sup>-8</sup> |  |  | F (2, 27) = 14.34, p = 5.7 x 10<sup>-5</sup> |  |  |
|  | Bonferroni post-hoc test | (C) > (A), (B); (A) □ (B) |  |  | (C) > (A), (B); (A) □ (B) |  |  |
| Experiment 2 | Colony Count | 7.1 ± 2.5 | 4.9 ± 1.4 | 21.9 ± 3.3 | 0.8 ± 0.6 | 0.2 ± 0.4 | 2.0 ± 1.6 |
|  | One-way ANOVA | F (2, 27) = 135.21, p = 8.6 x 10<sup>-15</sup> |  |  | F (2, 27) = 8.34, p = 1.5 x 10<sup>-3</sup> |  |  |
|  | Bonferroni post-hoc test | (C) > (A), (B); (A) □ (B) |  |  | (C) □ (A); (A) □ (B); (C) > (B) |  |  |
| Experiment 3 | Colony Count | 3.7 ± 2.1 | 1.8 ± 0.9 | 11.9 ± 3.3 | 0.6 ± 0.7 | 0.3 ± 0.5 | 1.7 ± 0.9 |
|  | One-way ANOVA | F (2, 27) = 53.79, p = 3.8 x 10<sup>-10</sup> |  |  | F (2, 27) = 10.05, p = 5.5 x 10<sup>-4</sup> |  |  |
|  | Bonferroni post-hoc test | (C) > (A), (B); (A) □ (B) |  |  | (C) > (A), (B); (A) □ (B) |  |  |

Fstatistic and p-value calculated from One-way ANOVA and interpretation of Bonferroni post-hoc tests for the different comparable groups are represented.

**Table 3(b).**
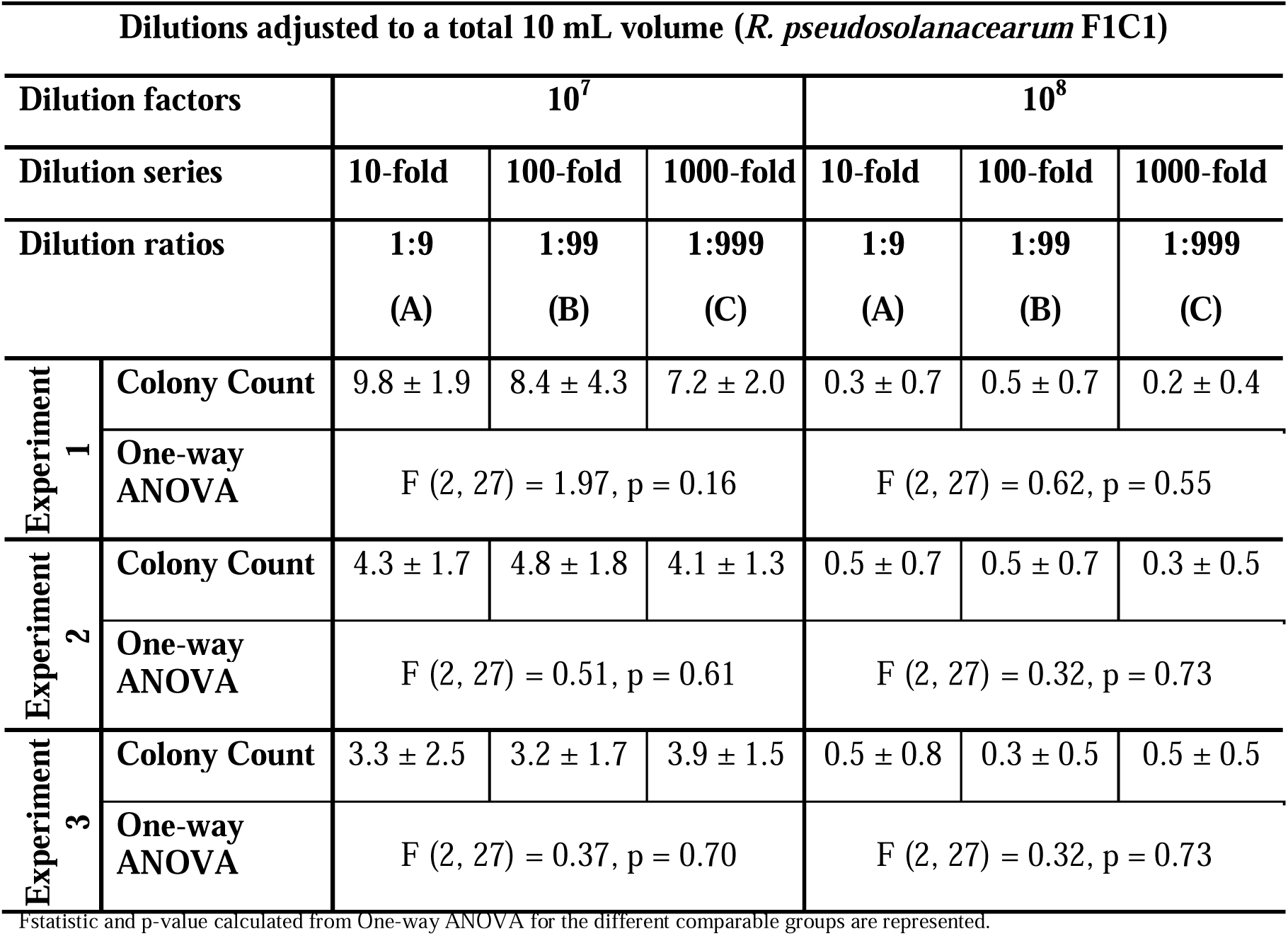
Average colony numbers (along with standard deviations) of *R. pseudosolanacearum* F1C1 obtained from different dilution series performed to a final 10 mL volume.

| Dilutions adjusted to a total 10 mL volume (*R. pseudosolanacearum* F1C1) |  |  |  |  |  |  |  |
| --- | --- | --- | --- | --- | --- | --- | --- |
| Dilution factors | | $10^7$ | | | $10^8$ | | |
| Dilution series |  | 10-fold | 100-fold | 1000-fold | 10-fold | 100-fold | 1000-fold |
| Dilution ratios |  | 1:9<br>(A) | 1:99<br>(B) | 1:999<br>(C) | 1:9<br>(A) | 1:99<br>(B) | 1:999<br>(C) |
| Experiment 1 | Colony Count | 9.8 ± 1.9 | 8.4 ± 4.3 | 7.2 ± 2.0 | 0.3 ± 0.7 | 0.5 ± 0.7 | 0.2 ± 0.4 |
|  | One-way ANOVA | F (2, 27) = 1.97, p = 0.16 |  |  | F (2, 27) = 0.62, p = 0.55 |  |  |
| Experiment 2 | Colony Count | 4.3 ± 1.7 | 4.8 ± 1.8 | 4.1 ± 1.3 | 0.5 ± 0.7 | 0.5 ± 0.7 | 0.3 ± 0.5 |
|  | One-way ANOVA | F (2, 27) = 0.51, p = 0.61 |  |  | F (2, 27) = 0.32, p = 0.73 |  |  |
| Experiment 3 | Colony Count | 3.3 ± 2.5 | 3.2 ± 1.7 | 3.9 ± 1.5 | 0.5 ± 0.8 | 0.3 ± 0.5 | 0.5 ± 0.5 |
|  | One-way ANOVA | F (2, 27) = 0.37, p = 0.70 |  |  | F (2, 27) = 0.32, p = 0.73 |  |  |

Fstatistic and p-value calculated from One-way ANOVA for the different comparable groups are represented.

### 3.4. Bacterial enumeration is indifferent to the dilution volume used in serial dilution

To observe if dilution volume used in serial dilution has an influence on bacterial enumeration, each of the 10-, 100- and 1000-fold dilution series were performed from a saturated *E. coli* culture using different diluent volumes to reach the final volume of 1 mL and 10 mL respectively. CFU observed with the plating of 10^6^ and 10^7^ fold dilutions obtained from each of the three dilution series adjusted to the different dilution volumes are represented in Table 4 and a representative result is shown in Figure 5. With 10^6^ fold dilution, a 10-fold series yielded 21.9 ± 3.0 and 18.1 ± 3.4, a 100-fold series yielded 19.7 ± 3.3 and 19.1 ± 3.4, and a 1000-fold series yielded 46.9 ± 5.3 and 19.9 ± 2.6 average numbers of colonies for the final 1 mL and 10 mL dilution volumes respectively. Similarly, with 10^7^ fold dilution, a 10-fold series yielded 2.6 ± 1.1 and 2.4 ± 1.3, a 100-fold series yielded 2.5 ± 1.3 and 1.9 ± 1.0, and a 1000-fold series yielded 5.6 ± 2.1 and 1.9 ± 1.3 average numbers of colonies for the final 1 mL and 10 mL dilution volumes respectively. Clearly, the colony numbers observed with the 1 mL and 10 mL dilution volumes were similar for most of the times for a 10-fold and 100-fold series; however they were greatly different in all the cases for the 1000-fold series. This suggests that the use of two dilution volumes brought no contrasting difference in the bacterial density being passaged in the 10-fold and 100-fold dilution series. However, the passaging in 1000-fold series was contrastingly different in the two diluent volumes used. Here, the 1 μL passaging (in 1 mL dilution volume) produced significantly higher number of colonies than the passaging of 10 μL (in 10 mL dilution volume); the latter yielding comparable colonies to that of the 10-fold and 100-fold series performed in either dilution volumes. It correlates to our previous understanding of the complexities associated with 1 μL pipetting. Overall, this suggests that the dilution volume used in serial dilution doesn’t affect the accuracy of bacterial enumeration, given that 1 μL pipetting is avoided.

**Figure 5.**
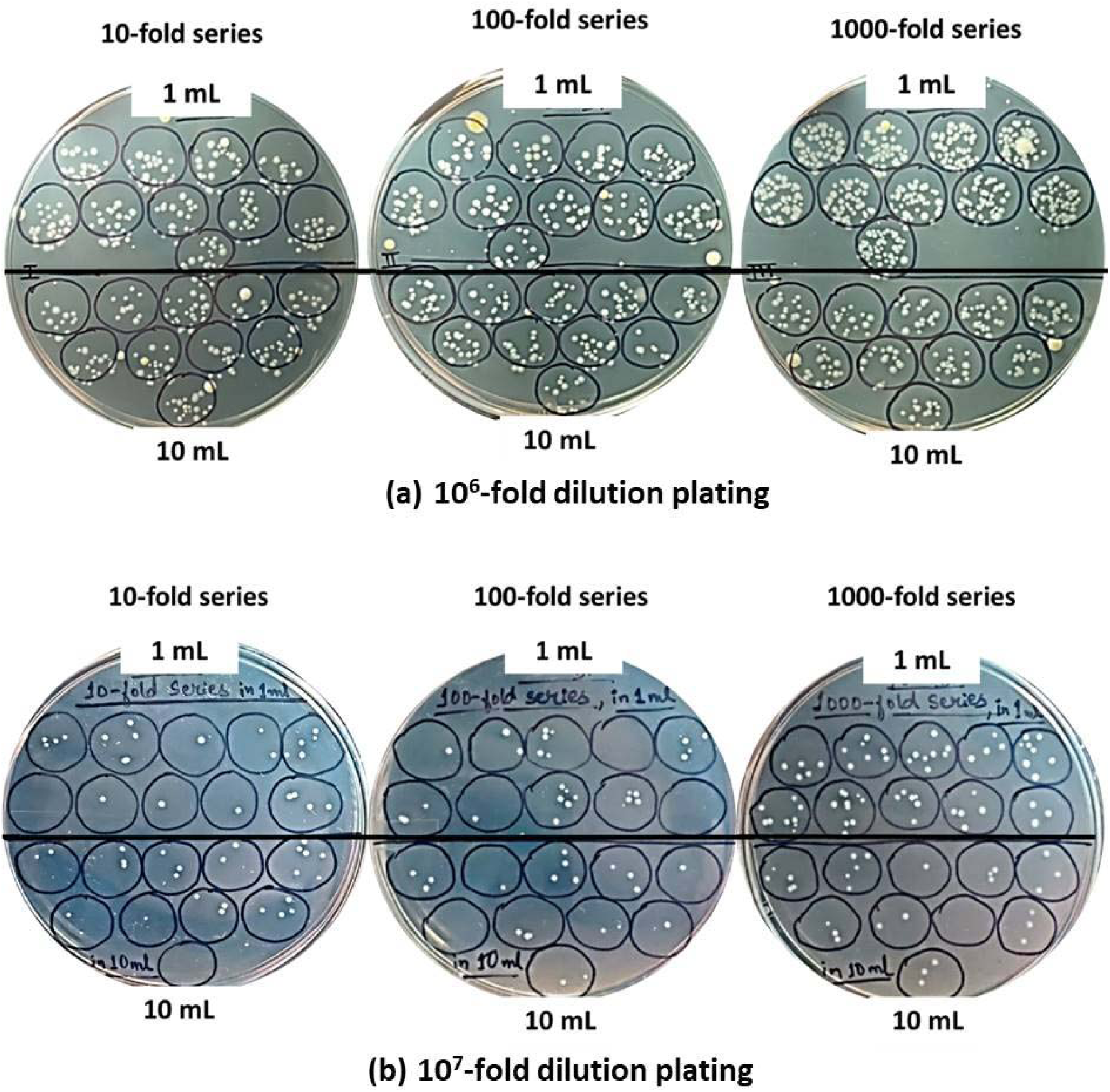
Comparison of colony numbers of *E. coli* generated from same culture but attained using two different diluent volumes of serial dilution in order to adjust to 1 mL and 10 mL volumes. Plating at (a) 10^6^- and (b) 10^7^-fold dilution was done. Each petri-plate corresponds to a specific dilution series where comparison is made between upper and lower halves containing colonies from 1 mL and 10 mL dilution volumes respectively. In each of the case, colony numbers observed with these dilution volumes appears similar for 10-fold and 100-fold series. However, a contrasting difference was observed with 1000-fold series, where a clearly greater number of colonies were observed in the case of 1 mL dilution volume

**Table 4.**
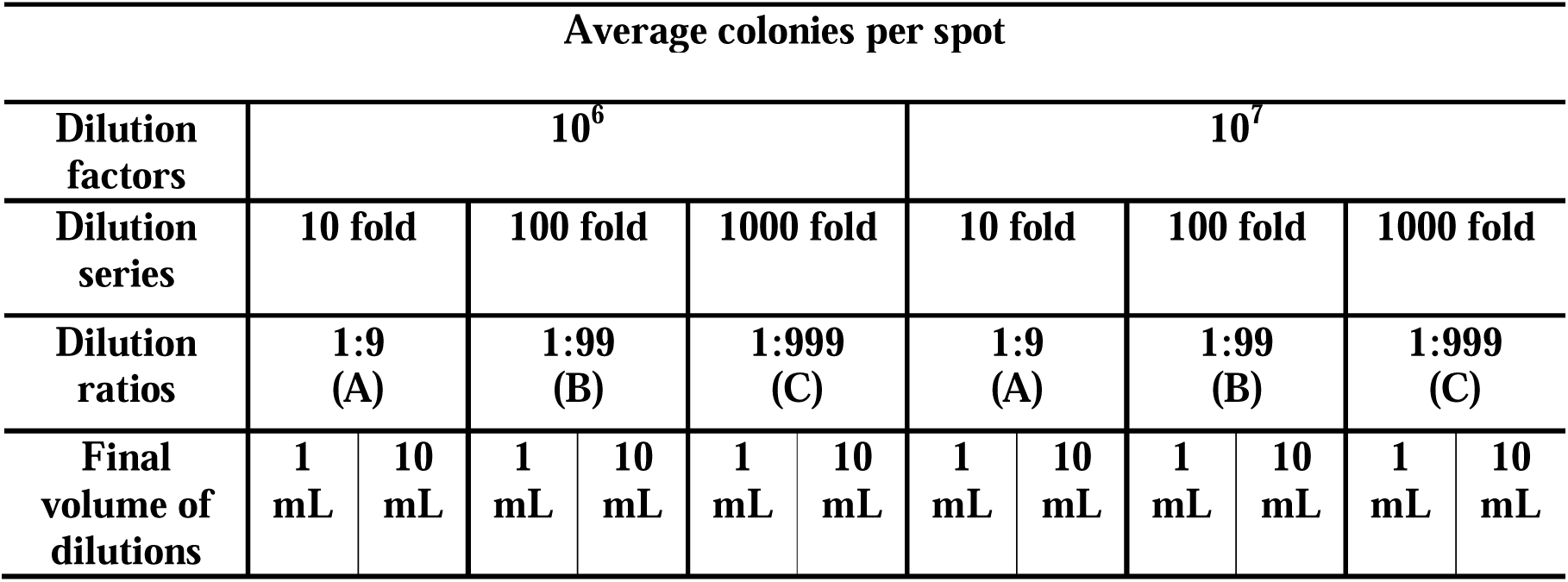
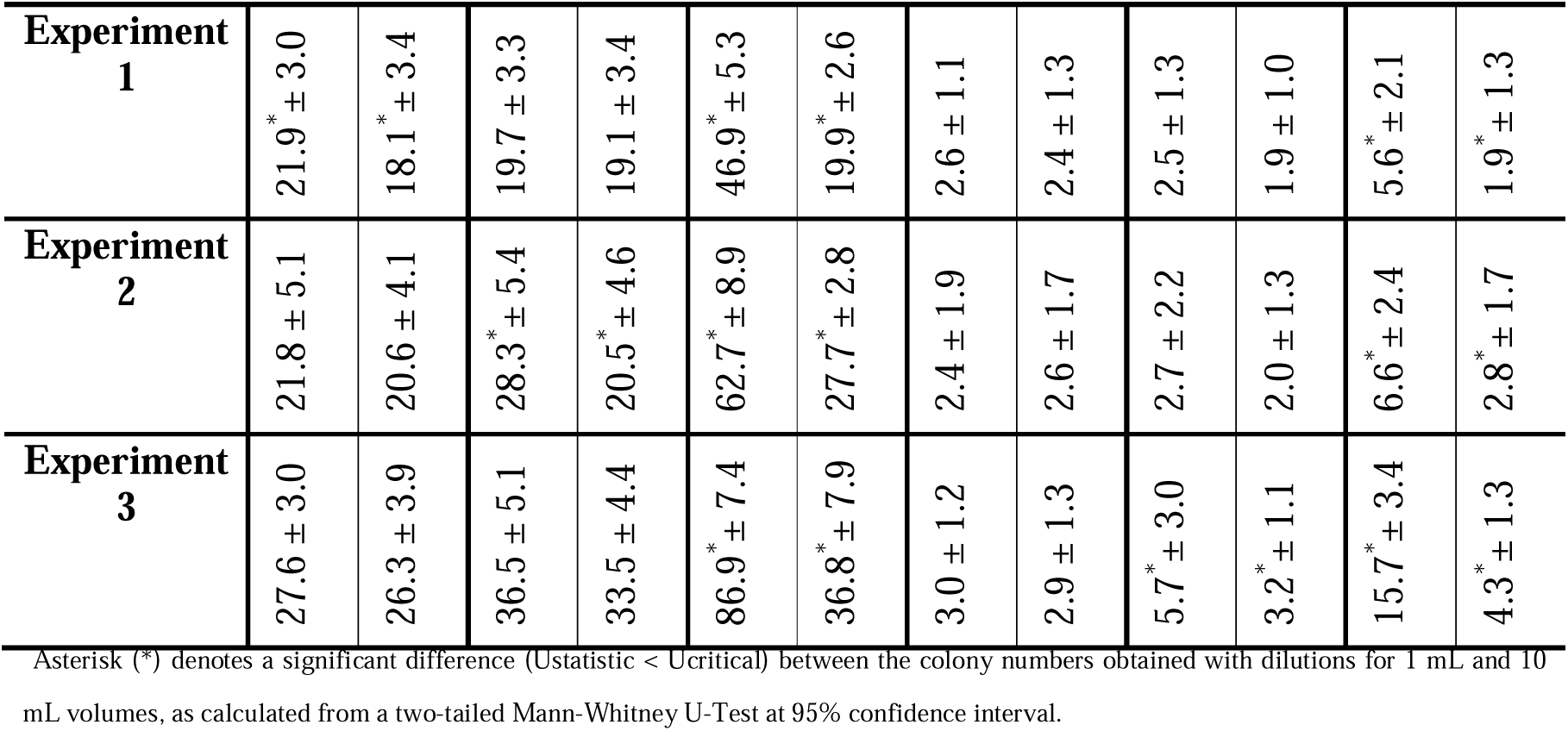
Comparison of the colony numbers (along with standard deviations) of *E. coli* obtained from each of the dilution series performed for 1 mL and 10 mL dilution volume, generated from same culture.

| Experiment<br>1 | 21.9<sup>*</sup> ± 3.0 | 18.1<sup>*</sup> ± 3.4 | 19.7 ± 3.3 | 19.1 ± 3.4 | 46.9<sup>*</sup> ± 5.3 | 19.9<sup>*</sup> ± 2.6 | 2.6 ± 1.1 | 2.4 ± 1.3 | 2.5 ± 1.3 | 1.9 ± 1.0 | 5.6<sup>*</sup> ± 2.1 | 1.9<sup>*</sup> ± 1.3 |
| --- | --- | --- | --- | --- | --- | --- | --- | --- | --- | --- | --- | --- |
| Experiment<br>2 | 21.8 ± 5.1 | 20.6 ± 4.1 | 28.3<sup>*</sup> ± 5.4 | 20.5<sup>*</sup> ± 4.6 | 62.7<sup>*</sup> ± 8.9 | 27.7<sup>*</sup> ± 2.8 | 2.4 ± 1.9 | 2.6 ± 1.7 | 2.7 ± 2.2 | 2.0 ± 1.3 | 6.6<sup>*</sup> ± 2.4 | 2.8<sup>*</sup> ± 1.7 |
| Experiment<br>3 | 27.6 ± 3.0 | 26.3 ± 3.9 | 36.5 ± 5.1 | 33.5 ± 4.4 | 86.9<sup>*</sup> ± 7.4 | 36.8<sup>*</sup> ± 7.9 | 3.0 ± 1.2 | 2.9 ± 1.3 | 5.7<sup>*</sup> ± 3.0 | 3.2<sup>*</sup> ± 1.1 | 15.7<sup>*</sup> ± 3.4 | 4.3<sup>*</sup> ± 1.3 |

Asterisk (\*) denotes a significant difference ( $U_{\text{statistic}} < U_{\text{critical}}$ ) between the colony numbers obtained with dilutions for 1 mL and 10 mL volumes, as calculated from a two-tailed Mann-Whitney U-Test at 95% confidence interval.

### 3.5 Bacterial enumeration is indifferent to the cell-density of the diluted suspension from which sampling is being carried out

To study whether the sampling error in bacterial enumeration is related with the cell-density of the diluted suspension from which sampling is being carried out, the 2-fold, 4-fold and 8-fold diluted suspensions were secondarily generated from the 10^6^-fold diluted suspension and plated as mentioned. Cell-density is highest in the 2-fold diluted suspension followed by the 4-fold and lowest in the 8-fold diluted suspension. The colony counts obtained in each of these cases are represented in S. Table 2. In Trial 1, the mean colony count obtained in case of the 2-fold, 4-fold and 8-fold dilutions are 21.50, 10.85 and 5.55 respectively. The decreasing factor from 2-fold to 4-fold is ∼1.98; from 4-fold to 8-fold is ∼1.96. These are very close to ‘2’ as also reflected by the dilution fold. The same is observed with other two Trials as well. Although the CoV is always higher with the 2-fold dilution, followed by the 4-fold and 8-fold dilution, but that represents an obvious outcome in view of the average colony counts in these cases. However, as the colony count ratio clearly depicts the dilution fold, it indicates that the sampling error during the enumeration process is not related with the cell-density of the diluted suspensions used for sampling. Therefore, bacterial enumeration is indifferent to the cell-density of the diluted suspension.

### 3.6 Bacterial enumeration is indifferent to the number of dilution passages

For this, the colony counts with the single-passage generated 4-fold and 8-fold diluted suspensions were compared with those of the 4-fold and 8-fold diluted suspensions obtained *via* the 2-fold series in two and three respective passages. The colony count data are presented in S. Table 2. In Trial 1, the average colony count with the single-passage generated 4-fold and two passages generated 4-fold diluted suspensions were observed to be 10.85 and 10.45 respectively. Similarly, the average colony count with the single-passage generated 8-fold and three passage generated 8-fold diluted suspensions were observed to be 5.55 and 5.75 respectively. The colony counts proved to be statistically similar as U_stat_>U_critical_ in each of the cases in the two-tailed Mann-Whitney U Test. The same was observed in two other Trials. This suggested that bacterial enumeration is indifferent to the number of serial dilution passages.

## 4. Discussion

In view of the Poisson or random nature of bacterial distribution in culture, sampling variability is inherent to the enumeration process of bacteria by dilution-plating method. Considering this, a sweet-spot has to be always sought to minimize the sampling error without compromising on the accuracy of enumeration; and accordingly, the suitable practices have to be selected. Sampling error during the dilution-plating of a randomly distributed population is attributable to several practices. Firstly, it inversely relates to the size of the population being sampled; such that a smaller sample volume yields greater variability than a larger sample volume. This presents a natural outcome of the random distribution. Secondly, the sampling error is also known to arise and increase progressively during the dilution process in the theoretical calculations of Ben-David and Davidson, 2014, where ‘*SD*/*mean*’ (CoV) has been used to refer to the relative sampling error. However, the empirical reports establishing the relation of sampling error with density of the suspension being sampled are lacking. Such information would prove useful in discriminating the choice of using a higher or lower dilution accordingly for enumeration.

In the current study, we first demonstrated that a higher sample volume is a better representation of the bacterial population size than a lower sample volume, evident from the values of CoV recorded (Table 1). In this regard, it was also interesting to observe that passaging different sample volumes in serial dilution stood statistically similar with regards to enumeration. For example, in 1 mL dilution volume, passaging either 100 µL or 10 µL or 1 µL (in the respective 10-, 100- and 1000-fold dilution series) produced similar cell counts with the exception of 1 µL passaging, which was found to produce more variable results. Notably, upon scaling up the volume 10-times, (in 10 mL dilution volume), passaging of 1 mL, 100 µL and 10 µL all produced similar cell counts suggesting that bacterial enumeration is indifferent to the passaged sample volume, given the passaging is done properly. 1 µL passaging might require a more sophisticated handling and has resulted in the observed inconsistencies. Considering it, passaging of a very small sample volume like 1 µL shall be avoided in the enumeration process. In the second part of the study, the cell count observed with the 2-fold, 4-fold and 8-fold diluted suspensions were found to decrease by the factor very close to 2, which is the well-expected outcome considering the dilution-folds. This again suggested that the enumeration that would be produced by passaging of 500 µL or 250 µL or 125 µL in 1 mL dilution volume (in the respective 2-, 4- and 8-fold dilution) would all stand comparable. These evidences also revealed that bacterial enumeration and the sampling error associated with bacterial sampling is indifferent to the volume passaged in serial dilution, even though a higher sample volume is a better estimate of the population size in context of the Poisson sampling.

It was also observed that the sampling error is indifferent towards the number of passages in a dilution series, evident from the instance that the 4-fold and 8-fold diluted suspension when attained in a single step produced similar cell counts to those attained in two and three respective steps *via* the 2-fold series. Same was also evident from the cell counts produced by the 10^6^-fold diluted suspensions, which when attained in six, three and two respective steps in the 10-, 100- and 1000-fold dilution series were found to produce similar cell counts.

Importantly, as the second part of the study suggested comparable enumerations by the 2-, 4- and 8-fold diluted suspensions, it shedded light on the density-independent nature of sampling error during bacterial sampling. As such, it also indicated that Poisson distribution in a bacterial culture remains unaffected by its cell density. The CoV although, was observed to be higher in case of the 8-fold series, followed by the 4-fold and 2-fold series, but that presented an obvious outcome; because in a Poisson distribution (where *mean* = *variance or √mean* = SD), CoV (given by *SD*/*mean*) naturally becomes 1/√*mean*. Observing a higher CoV in population with lower cell count is trivial in this context and shall not define the “sampling error” in the enumeration process as used by Ben-David and Davidson, 2014. In this view, our observation of the sampling error being indifferent to the bacterial cell density of the suspension was in contrast to the theoretical assumptions given by Ben-David and Davidson, 2014.

## 5. Conclusions

Putting altogether, our study concludes that for bacterial enumeration, only the dilution that facilitates colony counting is critical; bacterial enumeration is regardless of the way in which the dilutions are achieved. On that note, achieving a 10^7^-fold diluted suspension in 1 mL dilution volume by the classic method of performing seven 100 µL passages is equally preferable as: (i) achieving first a 10^6^-fold dilution in three 10 µL passages and then performing a 100 µL passage from it; or (ii) achieving first a 10^1^-fold dilution by one 100 µL passage and then performing three 10 µL passages from it. As in our study, bacterial enumeration by serial dilution has been proven to remain unaffected by the sample volume being passaged in serial dilution, the number of passages, cell-density of the diluted suspension and the dilution volume. Considering this, our study also encourages passaging a lower sample volume in serial dilution (given the pipettes are accurate), rather than a higher sample volume for bacterial enumeration, considering the time, cost and energy efficiency of the approach with no compromise on the accuracy of enumeration.

## Multidisciplinary Domains

This research covers the domains: (a) Microbiology (b) Statistics

## Acknowledgement

MJ is thankful to Tezpur University for the institutional fellowship as well as CSIR JRF fellowship. S Begum and SJG is thankful for the JRF fellowship from the DBT, GoI New Delhi grant (BT/PR41637/NER/95/1753/2021) awarded to SKR and also to Tezpur University for the institutional fellowship. S Bhuyan is thankful for the JRF/SRF fellowship from the UGC-NFSC, GoI, New Delhi. PRD is thankful for the CSIR JRF fellowship. LD is thankful for the DST-INSPIRE JRF/SRF fellowship. ND is thankful to DBT-supported Center for Bioinformatics and Computational Biology, Tezpur University (BT/PR40253/BTIS/137/52/2022) for the fellowship. SKR is thankful to his teachers: Prof. Ramesh V. Sonti and Prof. Anil K. Tripathi for teaching the fundaments of serial dilution.

## Declaration on AI Usage

The authors declare that the article has been prepared without the use of AI tools.

