## Supplementary File for "Theoretical expectations versus empirical observations in the bacterial enumeration process using serial dilution"

**Supplementary Materials**

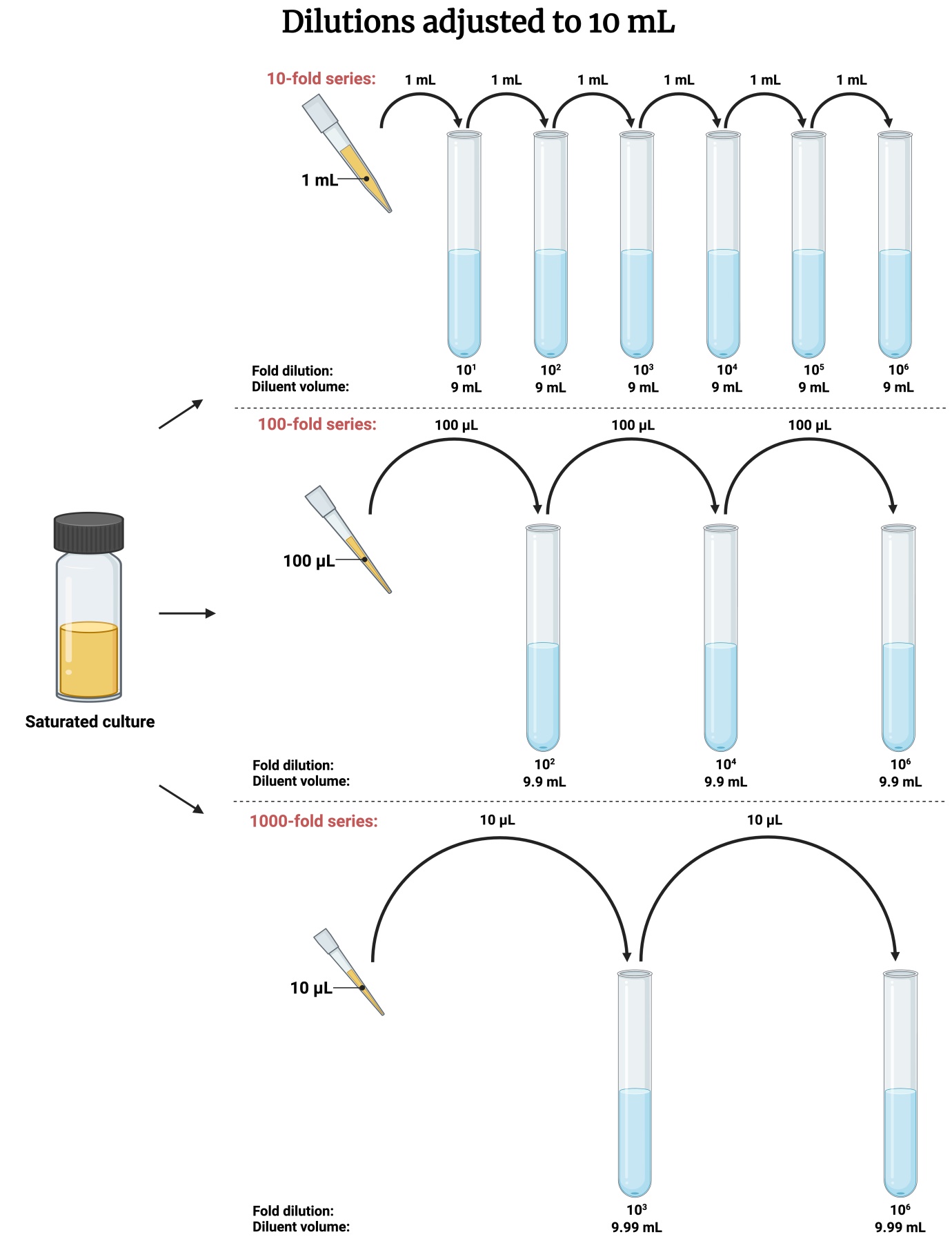

**S.Fig. 1** An illustration of the 10-, 100- and 1000-fold dilution series performed for 10 mL final volume. Created with Biorender.com

**(a)**

**(b)**
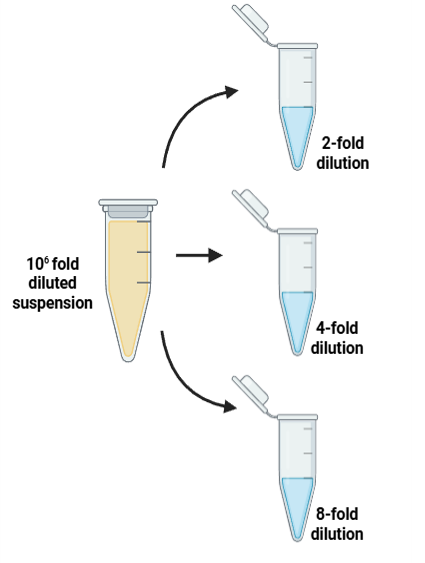

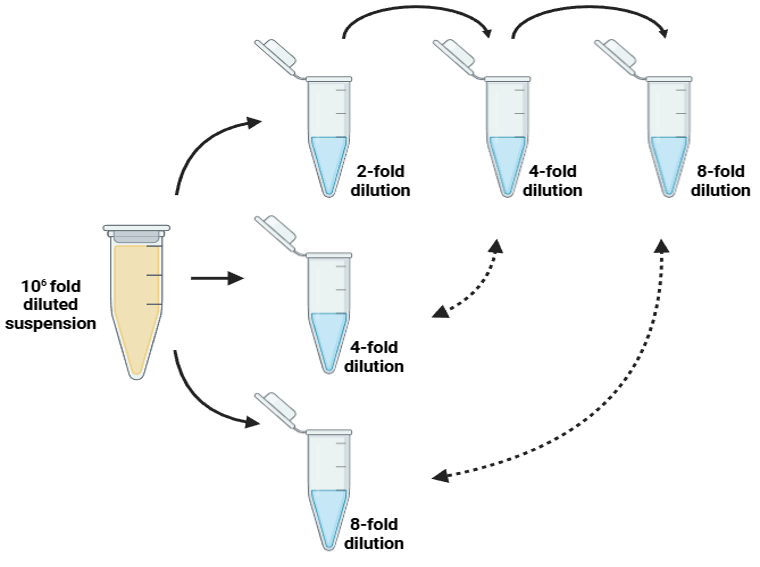

**S.Fig. 2** Simple illustrations of the different ways of attaining a dilution: **(a)** a single step-attained 2-fold, 4-fold and 8-fold dilutions from an already attained 10^6^-fold diluted suspension of a saturated *E. coli* culture **(b)** a two and three step attained 4-fold and 8-fold dilutions respectively besides the single step-attained dilutions

**S.Table 1.** Colony numbers obtained in different plating volumes of the two different bacterial cell suspensions.

|  | **Colony count data of 10 spots** | | | | | | | | | | |
| --- | --- | --- | --- | --- | --- | --- | --- | --- | --- | --- | --- |
|  | **Bacterial samples** | ***E. coli* DH5α** | | | | | | ***R. pseudosolanacearum* F1C1** | | | |
|  | **Dilution factors** | **10^6^** | | | | | | **10^7^** | | | |
| **Experiment 1** | **Spotting volume** | **5 μL** | | **10 μL** | | **15 μL** | **20 μL** | **5 μL** | **10 μL** | **15 μL** | **20 μL** |
|  |  | 7 | | 19 | | 36 | 38 | 0 | 5 | 2 | 6 |
|  |  | 7 | | 20 | | 32 | 37 | 1 | 1 | 2 | 8 |
|  |  | 12 | | 19 | | 27 | 46 | 1 | 3 | 4 | 7 |
|  |  | 9 | | 25 | | 30 | 35 | 0 | 2 | 8 | 11 |
|  |  | 7 | | 16 | | 30 | 42 | 2 | 2 | 5 | 9 |
|  |  | 5 | | 17 | | 33 | 45 | 2 | 4 | 4 | 10 |
|  |  | 12 | | 15 | | 26 | 43 | 1 | 2 | 5 | 6 |
|  |  | 7 | | 18 | | 25 | 47 | 1 | 4 | 1 | 8 |
|  |  | 11 | | 16 | | 26 | 38 | 3 | 2 | 6 | 6 |
|  |  | 8 | | 17 | | 32 | 48 | 2 | 5 | 4 | 4 |
|  | **Mean** | 8.50 | | 18.20 | | 29.70 | 41.90 | 1.30 | 3.00 | 4.10 | 7.50 |
|  | **SD** | 2.42 | | 2.86 | | 3.62 | 4.63 | 0.95 | 1.41 | 2.08 | 2.12 |
|  | **CoV** | 0.28 | | 0.16 | | 0.12 | 0.11 | 0.73 | 0.47 | 0.51 | 0.28 |
|  | **CFU estimate** | 1.6x10^9^ | | 1.9x10^9^ | | 1.7x10^9^ | 1.7x10^9^ | 2.6x10^9^ | 3.0x10^9^ | 2.7x10^9^ | 3.8x10^9^ |
| **Experiment 2** | **Spotting volume** | **5 μL** | | **10 μL** | | **15 μL** | **20 μL** | **5 μL** | **10 μL** | **15 μL** | **20 μL** |
|  |  | 10 | | 16 | | 28 | 38 | 2 | 7 | 7 | 14 |
|  |  | 5 | | 20 | | 24 | 34 | 3 | 2 | 9 | 10 |
|  |  | 10 | | 14 | | 28 | 25 | 4 | 8 | 6 | 13 |
|  |  | 7 | | 16 | | 21 | 41 | 6 | 8 | 14 | 16 |
|  |  | 7 | | 21 | | 26 | 32 | 1 | 3 | 13 | 11 |
|  |  | 7 | | 23 | | 33 | 27 | 2 | 1 | 3 | 9 |
|  |  | 8 | | 22 | | 19 | 33 | 3 | 5 | 14 | 15 |
|  |  | 11 | | 24 | | 21 | 32 | 5 | 12 | 5 | 6 |
|  |  | 7 | | 18 | | 28 | 39 | 1 | 4 | 8 | 10 |
|  |  | 10 | | 20 | | 29 | 33 | 4 | 9 | 8 | 11 |
|  | **Mean** | 8.20 | | 19.40 | | 25.70 | 33.40 | 3.10 | 5.90 | 8.70 | 11.50 |
|  | **SD** | 1.93 | | 3.31 | | 4.37 | 5.02 | 1.66 | 3.48 | 3.83 | 3.03 |
|  | **CoV** | 0.24 | | 0.17 | | 0.17 | 0.15 | 0.54 | 0.59 | 0.44 | 0.26 |
|  | **CFU estimate** | 1.6x10^9^ | | 1.9x10^9^ | | 1.7x10^9^ | 1.7x10^9^ | 4x10^9^ | 3.2x10^9^ | 3.3x10^9^ | 2.7x10^9^ |
| **Experiment 3** | **Spotting volume** | **5 μL** | | **10 μL** | | **15 μL** | **20 μL** | **5 μL** | **10 μL** | **15 μL** | **20 μL** |
|  |  | 25 | | 40 | | 53 | 72 | 1 | 3 | 4 | 4 |
|  |  | 19 | | 37 | | 53 | 72 | 1 | 2 | 2 | 4 |
|  |  | 14 | | 38 | | 56 | 56 | 0 | 2 | 3 | 6 |
|  |  | 16 | | 31 | | 53 | 69 | 0 | 3 | 2 | 3 |
|  |  | 24 | | 33 | | 42 | 65 | 3 | 0 | 4 | 9 |
|  |  | 20 | | 43 | | 52 | 80 | 0 | 3 | 4 | 6 |
|  |  | 19 | | 30 | | 50 | 67 | 2 | 5 | 3 | 7 |
|  |  | 13 | | 42 | | 42 | 60 | 2 | 4 | 3 | 9 |
|  |  | 20 | | 39 | | 53 | 69 | 3 | 2 | 3 | 4 |
|  |  | 23 | | 36 | | 45 | 68 | 0 | 1 | 7 | 3 |
|  | **Mean** | 19.30 | | 36.90 | | 49.90 | 67.80 | 1.20 | 2.50 | 3.50 | 5.50 |
|  | **SD** | 4.06 | | 4.43 | | 5.04 | 6.63 | 1.23 | 1.43 | 1.43 | 2.27 |
|  | **CoV** | 0.21 | | 0.12 | | 0.10 | 0.10 | 1.02 | 0.57 | 0.41 | 0.41 |
|  | **CFU estimate** | 3.9x10^9^ | | 3.7x10^9^ | | 3.3x10^9^ | 3.4x10^9^ | 4.8x10^9^ | 3.9x10^9^ | 4.2x10^9^ | 5.9x10^9^ |
| **Experiment 4** | **Spotting volume** | **5 μL** | | **10 μL** | | **15 μL** | **20 μL** | **5 μL** | **10 μL** | **15 μL** | **20 μL** |
|  |  | 4 | | 12 | | 15 | 22 | 1 | 1 | 0 | 5 |
|  |  | 10 | | 9 | | 27 | 23 | 2 | 2 | 2 | 3 |
|  |  | 8 | | 10 | | 17 | 27 | 1 | 1 | 2 | 4 |
|  |  | 8 | | 14 | | 31 | 28 | 2 | 4 | 3 | 5 |
|  |  | 5 | | 13 | | 21 | 31 | 3 | 1 | 7 | 3 |
|  |  | 4 | | 11 | | 14 | 20 | 1 | 1 | 4 | 6 |
|  |  | 7 | | 10 | | 12 | 22 | 0 | 5 | 5 | 3 |
|  |  | 9 | | 13 | | 22 | 28 | 1 | 0 | 5 | 7 |
|  |  | 7 | | 8 | | 21 | 29 | 2 | 2 | 3 | 5 |
|  |  | 3 | | 15 | | 20 | 20 | 0 | 4 | 2 | 5 |
|  | **Mean** | 6.50 | | 11.50 | | 20.00 | 25.00 | 1.30 | 2.10 | 3.30 | 4.60 |
|  | **SD** | 2.37 | | 2.27 | | 5.87 | 4.03 | 0.95 | 1.66 | 2.00 | 1.35 |
|  | **CoV** | 0.36 | | 0.20 | | 0.29 | 0.16 | 0.73 | 0.79 | 0.61 | 0.29 |
|  | **CFU estimate** | 1.3x10^9^ | | 1.2x10^9^ | | 1.3x10^9^ | 1.3x10^9^ | 4.8x10^9^ | 3.9x10^9^ | 4.2x10^9^ | 5.9x10^9^ |

^Standard deviation (SD), Coefficient of Variation (CoV) and CFU estimate calculated for parent culture in each of these cases are represented.^

**S.Table 2.** Colony counts of *E. coli* in 20 µL spots of the 2-fold, 4-fold and 8-fold diluted suspensions generated secondarily from an already attained 10^6^-fold diluted suspension and the 4-fold and 8-fold diluted suspensions attained *via* the 2-fold series from the 10^6^-fold diluted suspension.

| **Spot No.** | **Trial 1** | | | | | **Trial 2** | | | | | **Trial 3** | | | | |
| --- | --- | --- | --- | --- | --- | --- | --- | --- | --- | --- | --- | --- | --- | --- | --- |
|  | **Single Step** | | | ***via* 2-fold series** | | **Single Step** | | | ***via* 2-fold series** | | **Single Step** | | | ***via* 2-fold series** | |
|  | **2-fold** | **4-fold** | **8-fold** | **4-fold** | **8-fold** | **2-fold** | **4-fold** | **8-fold** | **4-fold** | **8-fold** | **2-fold** | **4-fold** | **8-fold** | **4-fold** | **8-fold** |
| **1** | 15 | 8 | 2 | 9 | 3 | 29 | 12 | 11 | 10 | 5 | 28 | 9 | 4 | 12 | 9 |
| **2** | 20 | 14 | 4 | 6 | 7 | 19 | 17 | 3 | 15 | 10 | 33 | 16 | 7 | 8 | 7 |
| **3** | 23 | 8 | 5 | 13 | 3 | 26 | 8 | 3 | 13 | 10 | 18 | 9 | 11 | 15 | 4 |
| **4** | 20 | 11 | 4 | 13 | 9 | 31 | 15 | 7 | 14 | 6 | 24 | 9 | 6 | 15 | 10 |
| **5** | 17 | 12 | 6 | 15 | 8 | 20 | 9 | 6 | 13 | 5 | 21 | 21 | 12 | 19 | 2 |
| **6** | 19 | 13 | 4 | 7 | 6 | 23 | 12 | 9 | 7 | 8 | 21 | 12 | 12 | 11 | 2 |
| **7** | 35 | 10 | 3 | 14 | 3 | 25 | 11 | 4 | 18 | 10 | 16 | 9 | 3 | 13 | 7 |
| **8** | 23 | 8 | 10 | 4 | 4 | 27 | 12 | 7 | 11 | 4 | 27 | 11 | 9 | 13 | 7 |
| **9** | 25 | 9 | 4 | 13 | 10 | 21 | 11 | 8 | 13 | 6 | 28 | 9 | 6 | 18 | 6 |
| **10** | 20 | 11 | 3 | 11 | 3 | 23 | 17 | 4 | 20 | 4 | 29 | 13 | 10 | 10 | 4 |
| **11** | 28 | 16 | 7 | 9 | 8 | 22 | 11 | 6 | 15 | 4 | 19 | 12 | 10 | 10 | 4 |
| **12** | 18 | 6 | 7 | 14 | 4 | 15 | 17 | 9 | 18 | 4 | 25 | 12 | 6 | 9 | 6 |
| **13** | 23 | 6 | 6 | 7 | 6 | 31 | 13 | 13 | 11 | 6 | 27 | 9 | 5 | 12 | 2 |
| **14** | 21 | 11 | 8 | 7 | 7 | 26 | 13 | 13 | 16 | 7 | 21 | 12 | 6 | 9 | 7 |
| **15** | 14 | 10 | 7 | 8 | 5 | 22 | 13 | 6 | 10 | 8 | 25 | 18 | 8 | 10 | 8 |
| **16** | 24 | 18 | 10 | 8 | 4 | 22 | 19 | 10 | 16 | 6 | 25 | 13 | 11 | 11 | 10 |
| **17** | 16 | 12 | 5 | 12 | 9 | 36 | 11 | 13 | 22 | 4 | 31 | 10 | 4 | 13 | 8 |
| **18** | 24 | 10 | 8 | 12 | 9 | 30 | 13 | 8 | 14 | 8 | 32 | 14 | 8 | 17 | 9 |
| **19** | 26 | 9 | 3 | 11 | 3 | 17 | 11 | 5 | 18 | 7 | 26 | 10 | 4 | 15 | 4 |
| **20** | 19 | 15 | 5 | 16 | 4 | 27 | 13 | 4 | 10 | 7 | 15 | 14 | 9 | 15 | 4 |
| **Mean** | 21.50 | 10.85 | 5.55 | 10.45 | 5.75 | 24.60 | 12.90 | 7.45 | 14.20 | 6.45 | 24.55 | 12.10 | 7.55 | 12.75 | 6.00 |
| **SD** | 4.90 | 3.18 | 2.31 | 3.36 | 2.45 | 5.22 | 2.83 | 3.28 | 3.81 | 2.06 | 5.15 | 3.29 | 2.86 | 3.13 | 2.62 |
| **CoV** | 0.23 | 0.29 | 0.42 | 0.32 | 0.43 | 0.21 | 0.22 | 0.44 | 0.27 | 0.32 | 0.21 | 0.27 | 0.38 | 0.25 | 0.44 |

^Mean colony count of the single step 4-fold diluted suspension was found statistically similar to the 4-fold diluted suspension obtained in two steps^ *^via^* ^the 2-fold series in each of the three trials; as Ustatistic > Ucritical in a two-tailed Mann-Whitney U-Test at 95% confidence interval. Similarly, the mean colony count of the single step 8-fold diluted suspension was found statistically similar to the 8-fold diluted suspension obtained in three steps^ *^via^* ^the 2-fold series in each of the three trials.^
